# Fast and accurate detection of metal resistance genes using MetHMMDb

**DOI:** 10.1101/2024.12.26.629440

**Authors:** Karol Ciuchcinski, Mikolaj Dziurzynski

## Abstract

Heavy metal pollution poses a major environmental challenge, with microbial resistance to heavy metals offering potential solutions through bioremediation. Additionally, the presence and diversity of microbial metal resistance genes (MMRGs) could contribute to an ecosystem’s ability to adapt and recover from heavy metal contamination by maintaining essential microbial functions and promoting the cycling of nutrients under stress conditions. Thus, MMRGs may serve not only as markers of contamination but also as indicators of an ecosystem’s self-purification capacity and resilience to environmental disturbances. Here we present MetHMMDB, a database containing 254 profile Hidden Markov Models representing 121 MMRGs. Unlike traditional sequence-based resources, MetHMMDB relies on HMMs to improve detection sensitivity and functional specificity across microbial communities. Created through iterative database searches, sequence clustering, structural prediction, and manual annotation, MetHMMDB emphasizes functional annotation rather than gene classification. The database outperforms sequence-based approaches, identifying over twice as many MMRGs in metagenomic datasets, including those from extreme environments. Analysis of agricultural soil revealed distinct resistance profiles correlating with soil quality. MetHMMDB advances our understanding of microbial adaptation to heavy metal contamination while supporting environmental management strategies through improved identification and characterization of metal resistance mechanisms.

Database URL: https://github.com/Haelmorn/MetHMMDB.

## Introduction

Heavy metal pollution, driven by industrialization and human activities, is a growing environmental challenge with severe consequences for ecosystems and biodiversity (1–3). The impact of heavy metal contamination varies significantly across different environmental contexts. In aquatic ecosystems, metals like mercury, lead, and cadmium accumulate in sediments and biomagnify through food chains, leading to widespread toxicity in fish populations and other aquatic organisms (4). Terrestrial ecosystems face similar challenges, with agricultural soils particularly affected by the accumulation of metals from fertilizers, pesticides, and atmospheric deposition (5). Urban environments show increasingly concerning levels of metal contamination, primarily from industrial emissions, vehicle exhaust, and waste disposal (6). Combined, these pollutants pose significant risks to human health through direct exposure and food chain contamination, leading to various chronic diseases and developmental disorders (3).

As heavy metal contamination continues to escalate, there is a need for effective solutions for monitoring and mitigating these pollutants. One promising avenue lies in the study of microbial resistance mechanisms, as they are the primary mechanisms responsible for metals biotransformation in nature (7). Furthermore, they have the potential to be harnessed for bioremediation and environmental monitoring (8,9). Microbial metal resistance is mediated by a diverse set of genes and pathways that enable microorganisms to tolerate or detoxify metal ions in contaminated environments. Identifying these microbial metal resistance genes (MMRGs) is crucial for understanding microbial adaptation, ecological impacts, and for developing biotechnological applications. Unfortunately, many current detection technologies for identifying metal resistance genes face significant limitations. Traditional sequence similarity-based approaches, while valuable for identifying closely related genes, often fail to detect distant homologs or novel resistance mechanisms. Many studies rely on databases like BacMet (10–12), which, while being a valuable source of manually curated data, has not been updated since 2018 (13). Modern environmental studies, particularly those involving complex microbial communities, must shift from classical sequence-to-sequence comparison to more sophisticated approaches that can account for the vast diversity of resistance mechanisms, previously undescribed distant homologs and their functional contexts.

Profile Hidden Markov Models (HMMs) have emerged as a powerful alternative to traditional sequence similarity searches, offering improved sensitivity in detecting distant homologs (14,15). Unlike simple sequence alignment tools, HMMs can capture position-specific amino acid preferences and insertion/deletion patterns characteristic of protein families, making them particularly suitable for identifying diverse metal resistance genes (16). Recent advances in structural biology and machine learning have also highlighted the importance of incorporating structural information into gene detection methods, as protein structure often remains conserved even when sequences have diverged significantly (17). To address these challenges and leverage recent methodological advances, there is a clear need for a comprehensive, up-to-date resource that combines the sensitivity of HMM-based detection with detailed functional annotation. Here we present MetHMMDB, a comprehensive, manually annotated database of 254 profile Hidden Markov Models (HMMs), uniquely designed to detect and characterize bacterial metal resistance genes with enhanced sensitivity and functional specificity in diverse datasets.

## Materials and Methods

### Database Construction and Annotation

The initial dataset used to create MetHMMDB contained 374 metal resistance genes obtained previously (18). Those were searched against the SwissProt database (obtained 07.08.2024) using DIAMOND BLASTP (>90% identity/coverage), yielding 315 new sequences. The datasets were then combined and used as a query against the NCBI NR database (obtained 04.04.2024), producing 11,025 total proteins (19,20). Next, MMseqs2 clustering was performed via the *easy-cluster* subcommand, with identity and coverage cutoffs ranging from 0.5 to 0.99 to determine optimal parameters (21). Final value (0.90) was selected based on the number of singletons and mean cluster size. Following the clustering, non-singleton clusters were annotated using sequence-based (Pfam_scan.pl; v.37.1), and protein structure-based tools such as LocalColabFold, and FoldSeek (E-value ≤1e-3) (22–24). Finally, multiple sequence alignments were generated via MAFFT with the *–auto* flag (v7.525) and the profile HMM database was created using HMMER3 (v3.4) *hmmbuild* and *hmmpress* commands (25,26).

### Performance Evaluation

Database performance was benchmarked using a reference bacterial genome (*E. coli* K-12 substr. MG1655; GCF_000005845.2) and a metagenomic dataset (∼150,000 proteins). Search speed comparisons between hmmsearch and DIAMOND BLASTP were conducted using hyperfine with 20 CPU threads and 3 warm-up runs to reduce the impact of IO/caching on results (https://github.com/sharkdp/hyperfine). Search sensitivity was evaluated using environmental metagenomes (PRJNA689378, PRJEB22376) by comparing *hmmscan* against DIAMOND BLASTP searches of the source database. For both methods, results were filtered using E-value cutoff of 1e-5.

## Results and Discussion

The MetHMMDB database was constructed through a systematic, multi-step process starting with 374 experimentally confirmed metal resistance genes from the MetGeneDB database, previously described in our work (18)(Fig 1A). Those sequences served as queries in DIAMOND BLASTP searches against SwissProt, using stringent criteria (>90% identity and coverage), yielding 315 new sequences. The combined set of 681 sequences was then used to search the NCBI NR database, identifying 10,344 additional unique sequences and bringing the final dataset to 11,025 proteins. These sequences were then clustered using MMseqs2 with clustering parameters optimized through a gradient of identity and coverage values. A value of 0.90 was selected, balancing mean cluster size and singleton count (Fig 1B). This resulted in 341 non-singleton clusters with a mean size of 27.98 proteins. The largest clusters included sequences encoding genes *mdtC* (811 sequences, represented by protein HCR6247170.1), *mdtB* (632 sequences; P69340), *cutC* (242 sequences; 1X7I), *fieF* (211 sequences; WP_085449461.1), and *zupT* (195 sequences; Q32BU5).

**Figure 1.**
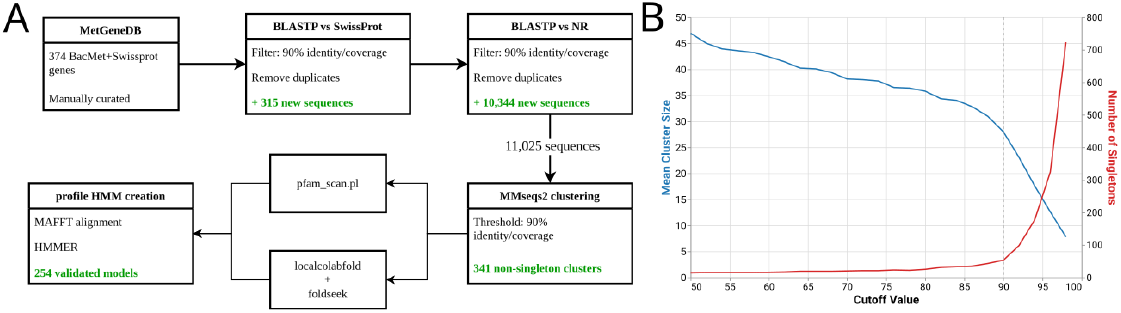
A) Database creation pipeline. Each box represents a separate step in the pipeline. B) Results of clustering parameters optimisation process. Coverage and identity value cutoffs are present on X-axis, while mean cluster size and number of singleton clusters can be seen on Y-axis.

For each non-singleton cluster, a comprehensive functional annotation using multiple complementary approaches was performed. The analysis combined domain detection using Pfam_scan.pl, protein structure prediction via LocalColabFold, and structural similarity searches against SwissProt using FoldSeek. Based on the resulting domain architectures and structural homologies to known SwissProt proteins, we determined the general function, metal specificity, and appropriate gene nomenclature for each cluster. This process identified 254 clusters associated with metal resistance mechanisms. Among these, copper-related clusters were most abundant (47), followed by clusters involved in arsenic and mercury resistance (32 each). For annotated clusters, multiple sequence alignments were generated using MAFFT and converted to profile HMMs using HMMER3. Each cluster was then assigned a descriptive name containing the metal specificity, general function, and gene name when possible.

To evaluate the practical utility of MetHMMDB, we compared its search speed to DIAMOND BLASTP. To do so, we used the *E. coli* K-12 substr. MG1655 genome (GCF_000005845.2, 4,298 protein sequences) and a subsampled metagenomic dataset (150,000 protein sequences). Both datasets were used as input for *hmmsearch* with MetHMMDB and *DIAMOND BLASTP* with MetGeneDB, and the speed of the search was measured using *hyperfine* (Table 1). Overall, the HMM-based approach was slower. Specifically, the *E. coli* genome was annotated in approximately 5.5s, compared to 0.85s of DIAMOND BLASTP approach. Furthermore, the difference increased as the size of the dataset grew, with *DIAMOND* being over 30x faster than *hmmsearch*.

**Table 1.** Performance comparison of protein vs HMM-based search.

| Dataset | DIAMOND BLASTP | hmmsearch | Speed Difference |
| --- | --- | --- | --- |
| *E. coli* (4,298 proteins) | 0.85s | 5.47s | 6.42× |
| Metagenome (150,000 proteins) | 1.53s | 52.88s | 34.43× |

Overall, we believe that while hmmsearch exhibited slower processing times than DIAMOND, its performance remained within acceptable limits for practical applications, annotating a complete genome in 5.5s and a metagenome in under a minute.

To evaluate resistance detection capabilities of the database, we analyzed a metagenome from a metal-rich Alpine stream (PRJNA689378) containing 215,000+ protein sequences (27). Using an E-value cutoff of 1e-5, hmmscan with MetHMMDB identified 3,846 metal resistance genes compared to 1,786 found by DIAMOND blastp with MetGeneDB. Both methods revealed similar distributions of resistance types, with *czcA*-like cobalt-zinc-cadmium resistance proteins and *fbpC*-like iron transporters being predominant. An interesting result can be observed for arsenic resistance, where the HMM-based approach demonstrated enhanced sensitivity. It not only successfully detected previously reported *Acr3* and *ArsACHM* arsenic resistance genes but also the presence of multiple copies of the *aioAB-*like arsenite oxidase - a finding not reported in the original study. A similar case was observed for silver resistance, where the original study only reported the presence of *silP* P-type ATPase. While the HMM-based results did not report the presence of genes designated as *silP*, closer investigation revealed that the same proteins were annotated as general silver and copper exporting ATPases, preserving the general function. Additionally use of MetHMMDB revealed the presence of *silABC* genes not reported in the original study. Despite those incongruencies, analysed methods showed substantial agreement, with 1,636 proteins (91.6% of blastp results) detected by both approaches. The HMM-based dataset missed primarily protein sequences with low sequence conservation (mean identity 50.1%, coverage 26.3%), including *acn* (aconitate hydratase), *nrsD* (MFS superfamily transporter), and *pitA* (inorganic phosphate transporter) genes.

Finally, we checked the performance of the database on agricultural soil metagenomes from BioProject PRJEB22376 (ERR2486626, ERR2486627) to evaluate its ability to differentiate healthy and unhealthy soil samples. The analysis revealed distinct metal resistance profiles between metagenomes, with multidrug *mdtBC* genes and iron transport related genes enriched in healthy soil, while nickel transporters and *fecE*-like iron transporters were more abundant in unhealthy soil (Supplementary Figure 1). Overall, the healthy soil exhibited a slightly increased diversity of metal resistance genes, with 195 unique HMMs found in 11,623 protein sequences, compared to 180 HMMs in 9,517 proteins in unhealthy soil. One of the most notable observed differences was seen in two *znuC* zinc transporter variants (Fig. 2). The first variant, more abundant in healthy soil, exhibits high similarity to proteins originating from environmental *Bacillus* strains. On the other hand, the *znuC* variant identified within unhealthy soil is most similar to various *Salmonella* strains, including pathogens such as *Salmonella typhi, Salmonella enterica* and *Salmonella typhimurium*. While it is possible that those differences may significantly contribute to the perceived soil quality, they can also be the result of many factors, such as usage of fertilizers [10.3389/fmicb.2019.00967].

**Figure 2.**
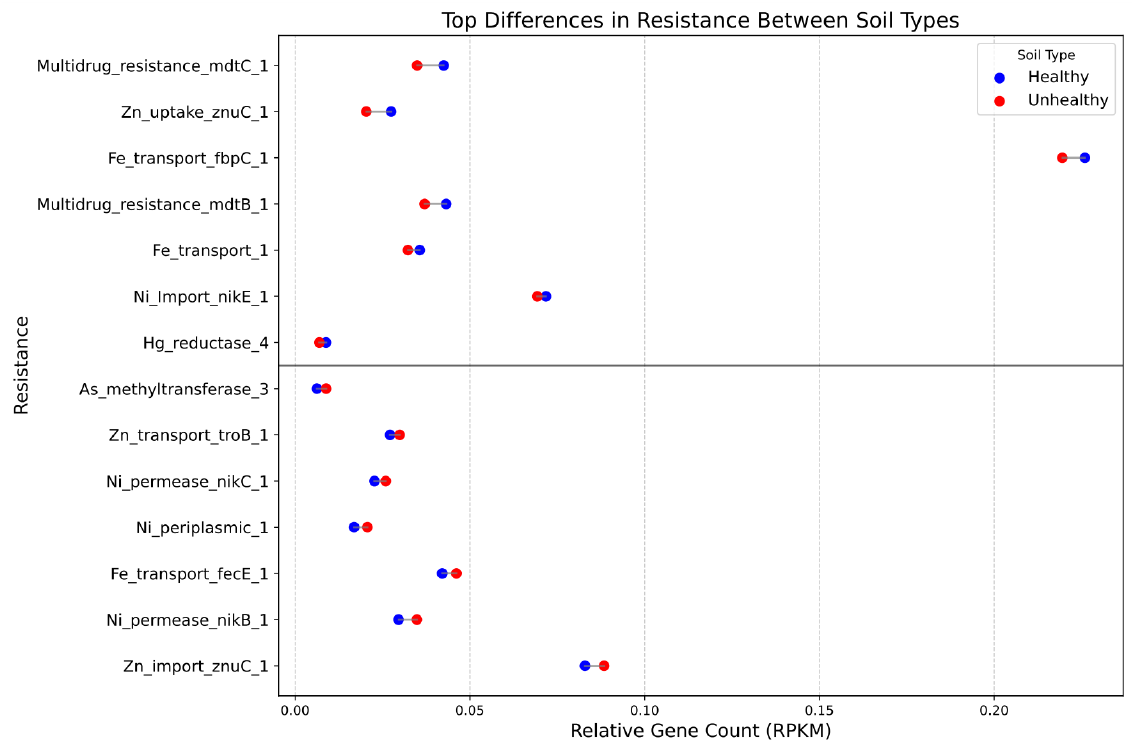
Comparison of the relative abundance of identified resistance mechanisms/genes between healthy and unhealthy soil samples. The X-axis represents the normalized proportion of gene counts (RPKM), indicating the relative frequency of identified genes: higher values to the right correspond to more genes detected, while lower values to the left correspond to fewer genes.

## Database availability

The database, including reference data and individual models, is available at https://github.com/Haelmorn/MetHMMDB. Hosting the database on GitHub ensures its persistence and broad accessibility. Given the rapidly expanding number of sequences available in public repositories used to build MetHMMDB, we plan to release updated versions of the database annually.

## Conclusion

In summary, MetHMMDB represents the first profile HMM-based database, specifically designed for MMRGs. By integrating sequence analysis, structural predictions, and manual curation, we developed 254 functionally annotated models that prioritize molecular function over traditional gene nomenclature. This approach improves identification accuracy and enables the detection of resistance determinants across diverse environmental contexts. The database’s utility was demonstrated through applications to both extreme environments and agricultural soils. Our analyses showed that MetHMMDB offers superior detection sensitivity and functional specificity compared to traditional sequence-based methods, identifying a broader range of MMRGs while providing ecological insights. Beyond its role in identifying resistance determinants, MetHMMDB has practical implications for understanding and managing environmental systems. The detection of MMRGs can be used to assess ecosystem resilience, offering insights into an environment’s ability to recover from anthropogenic or natural stressors. Additionally, the presence of HMRGs can serve as a marker of an ecosystem’s self-purification capacity, indicating its ability to mitigate contamination and maintain ecological stability. Though computationally more intensive, the database remains practical for large-scale analyses and is particularly well-suited for environmental monitoring and bioremediation research. With its focus on functional annotation rather than sequence similarity, MetHMMDB provides a robust tool for ecological and environmental studies, contributing to our understanding of ecosystem health and functionality

## Supplementary Data

**Supplementary Figure S1.**
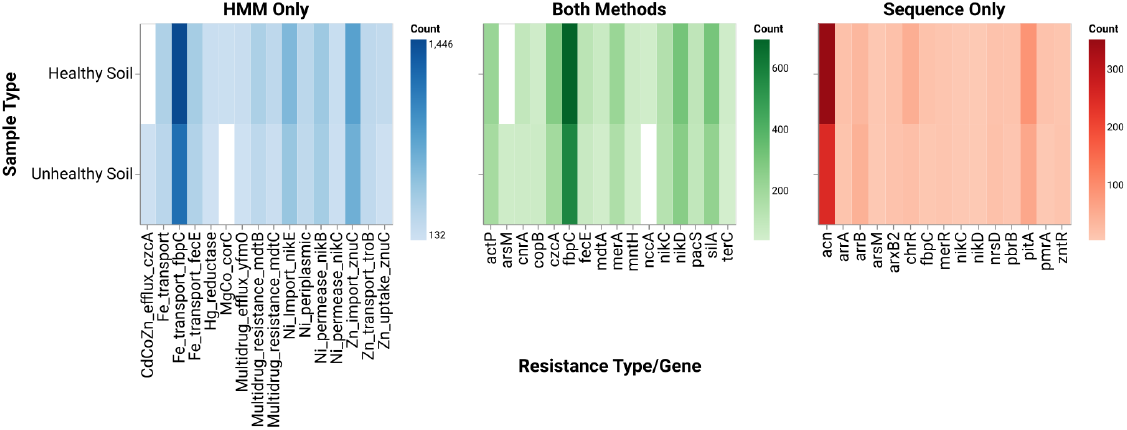
Comparative analysis of metagenome search results between healthy and unhealthy soil samples. The figure is organized in a 2×3 grid layout. Top row represents healthy soil samples, while the bottom row shows unhealthy soil samples. Left column displays sequences unique to Hidden Markov Model (HMM) search, middle column shows sequences common to both search methods, and right column indicates sequences unique to BLASTp search

## AUTHOR CONTRIBUTIONS

KC and MD conceptualized the study. KC performed data curation, formal analysis, validated and visualised the data. KC and MD conducted the investigations. KC wrote the original draft. KC and MD reviewed and edited the manuscript.

## REFERENCES

1. Das, S., Sultana, K.W., Ndhlala, A.R., et al. (2023) Heavy Metal Pollution in the Environment and Its Impact on Health: Exploring Green Technology for Remediation. Environ Health Insights, 17, 11786302231201259.

2. Darham, S., Zakaria, N.N., Zulkharnain, A., et al. (2023) Antarctic heavy metal pollution and remediation efforts: state of the art of research and scientific publications. Braz J Microbiol, 54, 2011–2026.

3. Briffa, J., Sinagra, E. and Blundell, R. (2020) Heavy metal pollution in the environment and their toxicological effects on humans. Heliyon, 6.

4. Zamora-Ledezma, C., Negrete-Bolagay, D., Figueroa, F., et al. (2021) Heavy metal water pollution: A fresh look about hazards, novel and conventional remediation methods. Environmental Technology & Innovation, 22, 101504.

5. Yang, Q., Li, Z., Lu, X., et al. (2018) A review of soil heavy metal pollution from industrial and agricultural regions in China: Pollution and risk assessment. Science of The Total Environment, 642, 690–700.

6. Pan, L., Wang, Y., Ma, J., et al. (2018) A review of heavy metal pollution levels and health risk assessment of urban soils in Chinese cities. Environ Sci Pollut Res, 25, 1055–1069.

7. Jing, R. and Kjellerup, B.V. (2018) Biogeochemical cycling of metals impacting by microbial mobilization and immobilization. Journal of Environmental Sciences, 66, 146–154.

8. Roy, R., Samanta, S., Pandit, S., et al. (2024) An Overview of Bacteria-Mediated Heavy Metal Bioremediation Strategies. Appl Biochem Biotechnol, 196, 1712–1751.

9. Jacob, J.M., Karthik, C., Saratale, R.G., et al. (2018) Biological approaches to tackle heavy metal pollution: A survey of literature. Journal of Environmental Management, 217, 56–70.

10. Li, L., Meng, D., Yin, H., et al. (2023) Genome-resolved metagenomics provides insights into the ecological roles of the keystone taxa in heavy-metal-contaminated soils. Front. Microbiol., 14.

11. Ekhlas, D., Cobo Díaz, J.F., Cabrera-Rubio, R., et al. (2023) Metagenomic comparison of the faecal and environmental resistome on Irish commercial pig farms with and without zinc oxide and antimicrobial usage. Animal Microbiome, 5, 62.

12. Balcha, E.S., Gómez, F., Gemeda, M.T., et al. (2023) Shotgun Metagenomics-Guided Prediction Reveals the Metal Tolerance and Antibiotic Resistance of Microbes in Poly-Extreme Environments in the Danakil Depression, Afar Region. Antibiotics, 12, 1697.

13. Pal, C., Bengtsson-Palme, J., Rensing, C., et al. (2014) BacMet: antibacterial biocide and metal resistance genes database. Nucleic acids research, 42, D737–43.

14. Feldgarden, M., Brover, V., Gonzalez-Escalona, N., et al. (2021) AMRFinderPlus and the Reference Gene Catalog facilitate examination of the genomic links among antimicrobial resistance, stress response, and virulence. Scientific reports, 11, 12728.

15. Reyes, A., Alves, J.M.P., Durham, A.M., et al. (2017) Use of profile hidden Markov models in viral discovery: current insights. AGG, 7, 29–45.

16. Keren, R., Méheust, R., Santini, J.M., et al. (2022) Global genomic analysis of microbial biotransformation of arsenic highlights the importance of arsenic methylation in environmental and human microbiomes. Computational and Structural Biotechnology Journal, 20, 559–572.

17. Koehler Leman, J., Szczerbiak, P., Renfrew, P.D., et al. (2023) Sequence-structure-function relationships in the microbial protein universe. Nat Commun, 14, 2351.

18. Dziurzynski, M., Gorecki, A., Decewicz, P., et al. (2022) Development of the LCPDb-MET database facilitating selection of PCR primers for the detection of metal metabolism and resistance genes in bacteria. Ecological Indicators, 145, 109606.

19. Buchfink, B., Reuter, K. and Drost, H.-G. (2021) Sensitive protein alignments at tree-of-life scale using DIAMOND. Nat Methods, 18, 366–368.

20. NCBI Resource Coordinators (2017) Database Resources of the National Center for Biotechnology Information. Nucleic acids research, 45, D12–D17.

21. Steinegger, M. and Söding, J. (2017) MMseqs2 enables sensitive protein sequence searching for the analysis of massive data sets. Nat Biotechnol, 35, 1026–1028.

22. Finn, R.D., Coggill, P., Eberhardt, R.Y., et al. (2016) The Pfam protein families database: towards a more sustainable future. Nucleic. Acids Res., 44, D279–D285.

23. Mirdita, M., Schütze, K., Moriwaki, Y., et al. (2022) ColabFold: making protein folding accessible to all. Nat Methods, 19, 679–682.

24. van Kempen, M., Kim, S.S., Tumescheit, C., et al. (2024) Fast and accurate protein structure search with Foldseek. Nat Biotechnol, 42, 243–246.

25. Katoh, K. and Standley, D.M. (2013) MAFFT Multiple Sequence Alignment Software Version 7: Improvements in Performance and Usability. Molecular Biology and Evolution, 30, 772–780.

26. Eddy, S.R. (2011) Accelerated profile HMM searches. PLoS Computational Biology, 7.

27. Buetti-Dinh, A., Ruinelli, M., Czerski, D., et al. (2021) Geochemical and metagenomics study of a metal-rich, green-turquoise-coloured stream in the southern Swiss Alps. PLOS ONE, 16, e0248877.

